# Multiplatform Biomarker Identification using a Data-driven Approach Enables Single-sample Classification

**DOI:** 10.1101/581686

**Authors:** Ling Zhang, Ishwor Thapa, Christian Haas, Dhundy Bastola

## Abstract

High-throughput gene expression profiles have allowed discovery of potential biomarkers enabling early diagnosis, prognosis and developing individualized treatment. However, it remains a challenge to identify a set of reliable and reproducible biomarkers across various gene expression platforms and laboratories for single sample diagnosis and prognosis. We address this need with our Data-Driven Reference (DDR) approach, which employs stably expressed housekeeping genes as references to eliminate platform-specific biases and non-biological variabilities. Our method identifies biomarkers with “built-in” features, and these features can be interpreted consistently regardless of profiling technology, which enable classification of single-sample independent of platforms. Validation with RNA-seq data of blood platelets shows that DDR achieves the superior performance in classification of six different tumor types as well as molecular target statuses (such as *MET* or *HER2*-positive, and mutant *KRAS, EGFR* or *PIK3CA*) with smaller sets of biomarkers. We demonstrate on the three microarray datasets that our method is capable of identifying robust biomarkers for subgrouping medulloblastoma samples with data perturbation due to different microarray platforms. In addition to identifying the majority of subgroup-specific biomarkers in Code-Set of nanoString, some potential new biomarkers for subgrouping medulloblastoma were detected by our method. Our results show that the DDR method contributes significantly to single-sample classification of disease and shed light on personalized medicine.

## Background

Identification of reliable and reproducible biomarkers can contribute to reveal patterns of disease heterogeneity. Recent advances in High-throughput sequencing (HTS) technology, such as microarray ^1,2^ and RNA-Seq ^3–5^ have enabled us to profile entire gene expression at low costs. The massive amounts of gene expression profile data generated by HTS have provided a great opportunity to identify reliable biomarkers which facilitate diagnosis, prognosis or treatment of patients. The technological biases across gene expression platforms and non-biological variabilities make it challenging to identify robust gene signature for cross-platform and cross-laboratory classification. Several techniques have been developed to eliminate platform-specific biases ^6–8^. However, these methods require multiple samples when processing transcriptome data, which is infeasible for analysis of biomarkers in samples obtained from single patient. Since gene expression data are high-dimensional data, an important research aim in analysis of transcription profiles is the discovery of small subset of biomarkers containing the most discriminant information, also known as feature selection ^9^, for accurate assignment of molecular subtype of disease. During the past years, numerous gene selection methods have been developed based on gene expression data and applied in disease classification. In general, the gene selection methods fall into four categories: filter methods ^10–12^, wrapper methods ^13–15^, hybrid methods ^16,17^ and embedded methods ^18,19^. However, there is a lack of feature selection methods designed to select robust features (or genes) which enable cross-platform classification of single disease sample.

To address these challenges, we present a Data-Driven Reference (DDR) approach to identify robust cross-platform gene signature for classification of single-sample from various platforms. Our DDR algorithm consists of three main steps: 1) the stably expressed housekeeping genes are employed as references to create a contingency table for each gene using given gene expression dataset; 2) Fisher’s exact tests are applied in contingency tables to identify differentially expressed genes (DEGs) as potential biomarkers between two conditions; 3) the categories which the expression levels of biomarkers fall into based on selected reference genes serve as input to the classifier. The categories generated by stably expressed reference genes represent “built-in” features that have a consistent interpretation across gene expression platforms and eliminate sample-specific biases. We illustrate DDR’s utility through various evaluations and comparisons with gene signatures identified by existing methods. We demonstrate that DDR method contributes significantly to identification of robust cross-platform gene signature for disease classification of single-patient to facilitate precision medicine.

## Results

### Identification of Potential Biomarkers in Various Expression Platforms

Differential expression analysis has been widely used to identify potential biomarkers for diagnosis and prognosis ^20^. Using DDR to identify discriminant genes between two conditions involves first two steps: constructing the contingency table for each gene from expression data based on selected reference genes, and then, using the Fisher’s exact test to determine if there is a significantly different expression for that gene between two groups (see **Methods** for details). For example, five reference genes (*STARD7-AS1, ZCCHC9, TIAL1, HNRNPH1*, and *EIF4G2*) from TCGA-BRCA RNA-Seq dataset were selected, so that log2-fold-changes between expressions of two consecutive reference genes were around 2 (Fig.1 and Supplementary Excel I). Gene expression heatmap was constructed to show the relative expression patterns of the top 20 (ranked based on FDR values) most significant DEGs in comparison between triple-negativebreast cancer (TNBC) and the other subtypes (Fig. S1A). Most of top 20 DEGs were down-regulated in TNBC samples compared with the other subtypes of breast cancer. TNBC has a poor prognosis compared with other types of breast cancer due to lack of therapeutic targets. In this study, we examined top 10 up-regulated genes (long non-coding RNA, LINC02188, was not included) (Fig. S1B), and at least six genes have been very recently (*BCL11A, FOXC1, CDCA7, PSAT1, UGT8*, and *GABRP*) experimentally validated for clinical or functional relevance in growth and metastasis of TNBC ^21–26^. Four other genes (*B3GNT5, PPP1R14C, RGMA*, and *HAPLN3*) were also computationally selected as signature genes in TNBC^27–29^. The DDR was also performed in LUAD RNA-Seq dataset from TCGA and DEGs were identified based on selected reference genes (Supplementary Excel II and Fig. S2). The method presented here can be applied as well to microarray expression data. Four reference genes (Fig. S3) were selected from expression microarray data (Accession: GSE62872), so that the differences between expressions of two consecutive reference genes were around 2. Then, the DEGs were identified by DDR and ranked by adjusted *p*-value (Supplementary Excel III).

**Figure 1:**
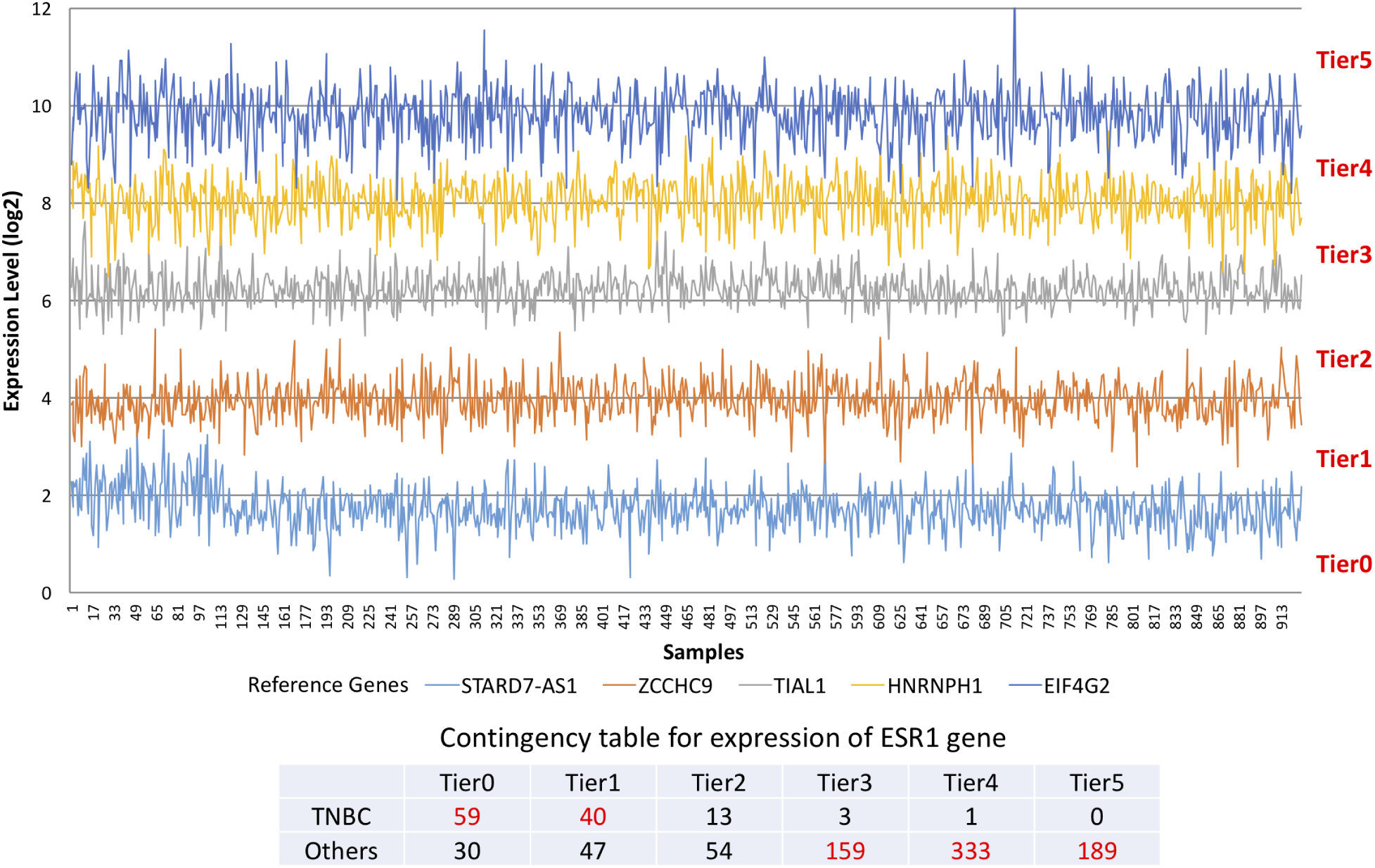
Expression levels of five data-driven reference genes from TCGA BRCA RNA-Seq samples and tiers classified based on reference genes (top). The contingency table of ESR1 expression based on expression levels of reference genes (bottom).

To assess DDR’s ability to detect DEGs, we compared it with the tools widely used in differential expression analysis in various platforms. The Fisher’s exact test is a nonparametric test in the sense that it does not assume that the RNA-Seq read counts or microarray expression data across samples are based on the theoretical probability distribution. On the contrary, current popular tools, such as *DESeq* ^30^, *DESeq2* ^31^ and *edgeR* ^32^, use a negative binomial distribution to model RNA-Seq read counts for assessing differential expression. Linear models for microarray (*limma*) ^33^ uses linear models based on empirical Bayes method to identify DEGs. To compare DDR with existing tools for analysis of RNA-Seq data, a gene was declared as significantly differentially expressed if FDR (or adjusted *p*-value) was less than 0.01 in *EdgeR* and *DESeq2* methods, or FDR (adjusted p-value) was less than 0.1 in *DESeq* and DDR methods. We measured the precision and recall of the identified DEGs using the DEGs from the datasets of 232 TCGA-BRCA samples (116 TNBC samples and randomly selected 116 other subtypes) and 118 TCGA-LUAD samples (59 normal tissue samples and randomly selected 59 LUAD samples) as the gold standard. The precision and recall values in both datasets for different methods and different numbers of samples per group are illustrated in Fig. 2. Two *EdgeR* methods reported high values for precision in both datasets across different sample sizes (Fig. 2A and Fig. 2D). *DESeq2* achieved high performance in precision similar to *EdgeR* methods in BRCA dataset, but showed a slight decrease in LUAD dataset (Fig. 2A and Fig. 2D). For DDR, the precision values remained relatively high in LUAD datasets when sample size was reduced and was slightly reduced in BRCA dataset (Fig. 2A and Fig. 2D). On the contrary, *DESeq* showed lower values for precision with respect to all other tools. The recall values rapidly decreased for all the tools when the number of samples per group was decreased (Fig. 2B and Fig. 2E). *DESeq2, EdgeR_GLM* and *EdgeR_EXACT* outperformed the other methods and *DESeq* was the worst-performing method in analysis for both datasets. DDR resulted in intermediate values of recall with respect to all other tools (Fig. 2B and Fig. 2E).

**Figure 2:**
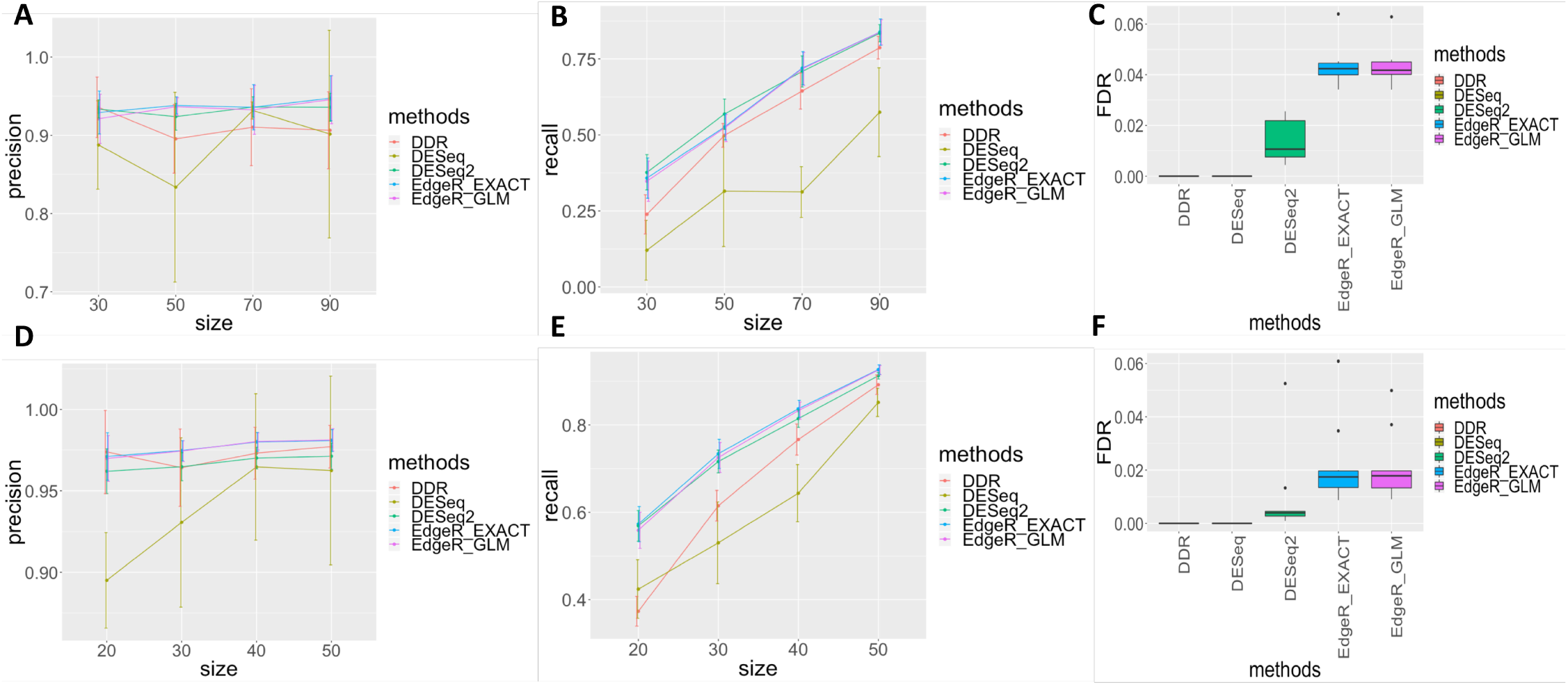
Precision and recall of methods in two TCGA RNA-Seq datasets: precision (**A**) and recall (**C**) in BRCA data and precision (**D**) and recall (**E**) in LUAD data. The false discovery rate (FDR) on the basis of mock comparisons generated using two datasets: 116 TNBC samples from TCGA-BRCA dataset (**C**) and 59 normal samples from LUAD (**F**).

To evaluate the false discovery rate (FDR) of the tools in analysis of RNA-Seq data, we generated mock comparisons from two datasets: the first consisted of 116 triple-negative breast cancer samples by randomly dividing the samples into two non-overlapping groups (58 samples for each group) and second consisted of 59 normal samples from TCGA-LUAD dataset by randomly dividing the samples into two non-overlapping groups (29 samples for one group and 30 samples for the other group). The median FDRs of both *EdgeR* methods were higher compared with the other methods in both datasets (Fig. 2C and Fig. 2F). *DESeq2* performed better than *EdgeR* methods and controlled the FDRs well (around 0.01 in BRCA dataset (Fig. 2C) and < 0.01 in LUAD dataset (Fig. 2F)). DDR and *DESeq* demonstrated extremely better control on false discovery rate compared with the other tools. It is essential to control false positives so that reliable and reproducible biomarkers can be identified.

Finally, we compared the overlaps of DEGs identified by the different methods through computing overlap coefficient (Szymkiewicz-Simpson coefficient) ^34^. The overlaps between the methods are listed in Table S4. In LUAD dataset, 80% of DEGs, identified using DDR (FDR < 0.1), coincided with DEGs identified using *EdgeR* (FDR < 0.01) or *DESeq2* (adjusted p-value < 0.01). The use of DDR and *DESeq2* (or *EdgeR*) algorithms achieved higher overlap rate in DEG results from BRCA dataset. DESeq generated DEG list overlapped poorly with that from DDR (< 52%) in both datasets. It is no surprise that the highest overlap percentages were observed between *DESeq* and *DESeq2* DEG lists or between *EdgeR_EXACT* and *EdgeR_GLM DEG* lists.

We benchmarked DDR approach against *limma* by using prostate cancer microarray data. Similarly as in analysis of RNA-Seq data, we used 240 samples (randomly selected 120 samples from each group) as the gold standard and measured the precision (Fig. S4A) and recall (Fig. S4B) for both methods in different sample size per group. *limma* performed better in term of precision. The recall values were systematically lower than precision values for DDR and *limma*. Fig. S4C shows that both DDR and *limma* appeared extremely conservative in controlling FDR in this analysis. DDR and *limma* DEG lists achieved 83% overlap with each other.

### Feature selection and cross-platform single-sample classification

In this section, we provided an example of using DDR to select signature genes between TNBC and other types of BRCA using TCGA RNA-Seq dataset, and use features of selected genes to classify BRCA samples from a different expression profiling platform (microarray). DDR was applied to TCGA-BRCA RNA-Seq dataset to identify a list of ranked DEGs (ranked by FDR) (Supplementary Excel I). Twenty top-ranked genes (Fig S1A) were sorted by expression distance (ED) and a set of top biomarkers was determined through comparing the classification performances of sets of top genes in terms of accuracy, recall and F1 score (Fig. S6A). Combination of FDR and ED for selection of signature genes enables not only identifying genes containing most discriminant information but also leading to more reproducible biomarkers. The smallest subset of 6 top genes (*ESR1, AGR2, FOXA1, AGR3, TFF3, MLPH*), achieving the best accuracy classification (Table 1A), was retained and their tier information (Supplementary Excel V) served as input to train classifiers. Here, we compared the performance of different classifiers from *Scikit-learn* ^35^ for TNBC classification using categorized expression of 6 signature genes as feature. From Fig. S6B, it can be seen that SVM achieved slightly better performance (Accuracy: 92%) though the other classifiers performed as well on classification task. Most of non-TNBC breast cancer samples were correctly predicted (Accuracy: 95%), whereas the proportion of mis-assigned TNBC samples was a little high (Table 1A). To evaluate the capacity of six selected signature genes and SVM classifier trained on TCGA-BRCA dataset in cross-platform classification of single-samples, the microarray dataset containing 5 TNBC samples and 14 non-TNBC samples was collected from GEO (Accession: GSE27447) ^36^. GSE27447 data (.CEL files) were normalized by *affy* package in R. When using tiered classifications of 6 genes selected above (Supplementary Excel V) based on reference genes from GSE27447 dataset as input to SVM classifier trained on TCGA-BRCA dataset, 4/5 (80%) and 12/14 (86%) were classified correctly to TNBC and non-TNBC, respectively (Table 1B). These examples demonstrate DDR’s ability to identify robust biomarkers for cross-platform classification of single patient.

**Table 1:**
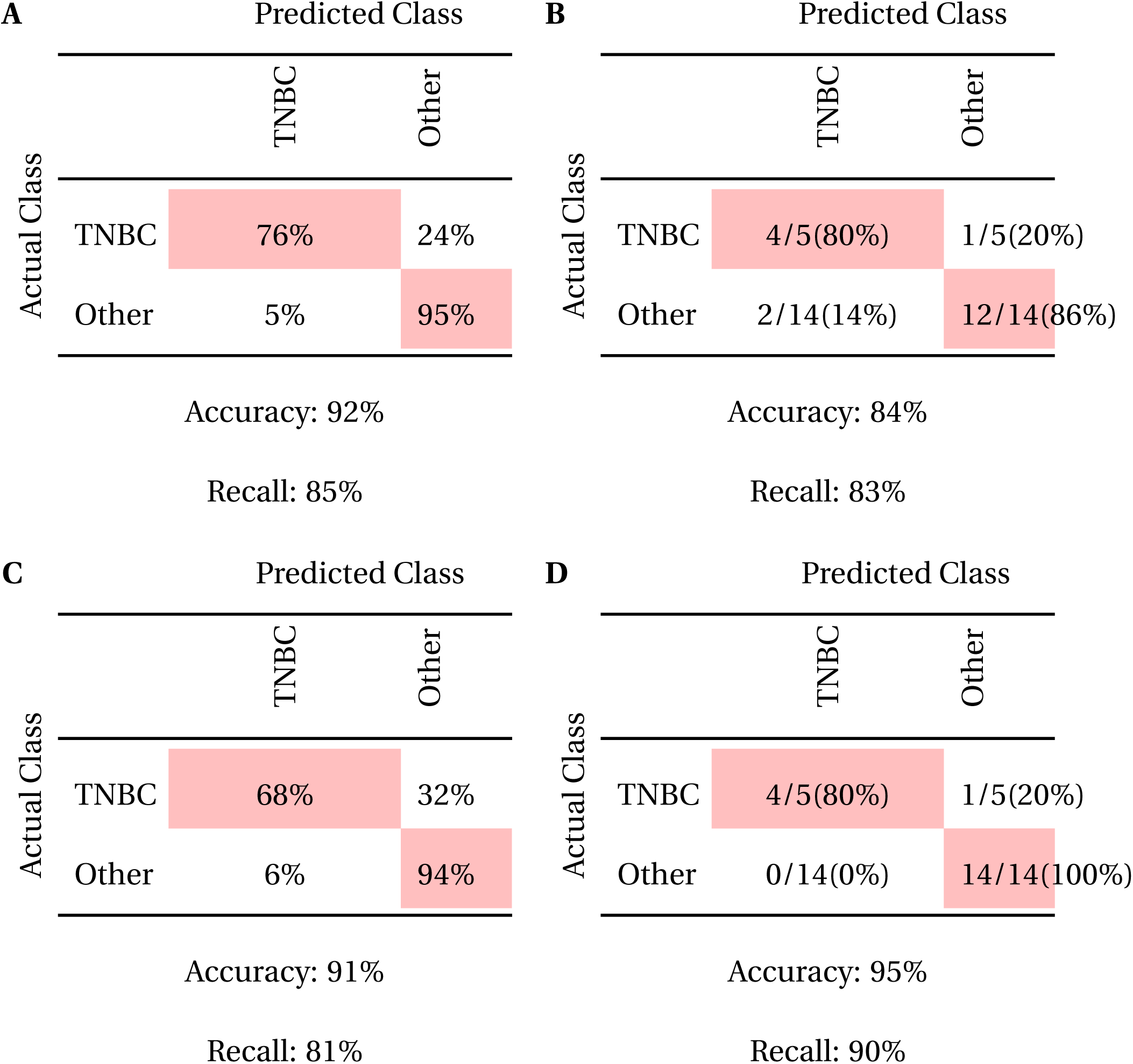
Classification Performance on (**A**) TCGA-BRCA RNA-Seq Dataset and (**B**) GSE27447 BRCA Microarray Test Dataset using 6 Signature Genes; Classification Performance on (**C**) TCGA-BRCA RNA-Seq Dataset and (**D**) GSE27447 BRCA Microarray Test Dataset using 2 Up-regulated genes

To examine subgrouping ability of up-regulated genes, two up-regulated (*BCL11A, B3GNT5*) genes from top 20 ranked genes were used as signature to classify TCGA-BRCA samples, which yielded an accuracy of 91%, and a recall of 81% (Table 1C). Subsequent validation using GSE27447 microarray dataset yield an accuracy of 95% and a recall of 90% when using SVM classifier trained on TCGA-BRCA dataset. Of note, 4 of the 5 TNBC samples and all non-TNBC samples were correctly classified (Table 1D). These examples demonstrate DDR’s ability to identify robust biomarkers for cross-platform classification of single patient.

### Identification and analysis of different cancer subtypes using RNA-Seq of Tumor-Educated Platelets

Molecular information in non-invasive liquid biopsy offers the promise of detection and classification of cancer subtypes ^38,39^. In this study, we employed RNA-Seq data of blood platelets to evaluate DDR in its ability to identify biomarkers for the classification of cancer subtypes and status of therapy-targeting genes. The platelets as liquid biopsy are capable of carrying RNA molecules from tumor tissues (educating), and serve as potential non-invasive biomarker source for detecting and monitoring cancers ^39,40^. These platelets are known as Tumor-Educated Platelets (TEP), of which the RNA profiles could be used to subgroup the cancers ^39^.

Here, we applied DDR to identify the subsets of discriminant genes from RNA-Seq data of TEP from GEO (Accession: GSE68086) ^39^ and employed SVM classifier on classification tasks. Furthermore, we compared the classification performance of our method with that of Best *et al.* (see **Discussion** for details) ^39^. To classify pan-cancer samples representing six tumor types (breast cancer (BRCA, n = 39), colorectal cancer (CRC, n = 42), glioblastoma (GBM, n = 40), non-small cell lung cancer (NSCLC, n = 60), hepatobiliary cancer (HBC, n = 14), and pancreatic cancer (PAAD, n = 35)) and healthy donors (HD, n = 55), the DDR was applied to each pairwise comparison among groups to identify DEGs, and then a small subsets of biomarkers were selected from DEG lists of pairwise comparisons based on adjusted *p*-values and ED (see Supplementary Excel VI for details). These subsets of biomarkers were merged and duplicate genes were removed to generate a list comprising 596 genes (Supplementary Excel VII) for pan-cancer classification. The tiered categorizations of 596 genes were used as input to multi-class One-versus-One (OvO) SVM classifier to yield overall accuracy of 72% (Table 2A). Similarly, for discriminating three different types of adenocarcinomas (CRC, PAAD and HBC) from gastro-intestinal tract, we selected 144 genes (Supplementary Excel VII) and performed OvO SVM classifier, yielding an overall accuracy of 80% (Table 2B). Best *et al.*^39^ also reported that RNA profiles from TEP could be used to discriminate tumor patients with different status of therapy-targeting oncogenes, such as *KRAS*-mut vs *KRAS*-wt in CRC, HBC, NSCLC, and PAAD patients, *EGFR*-mut vs *EGFR*-wt in NSCLC patients, *MET* + vs *MET* - in NSCLC patients, *PIK3CA*-mut vs *PIK3CA*-wt in BRCA, *HER2*+ vs *HER2*-in BRCA, as well as triple-negative breast cancer. The number of biomarkers (Supplementary Excel VI and VII) and classification accuracies were presented in Tables 3A-3I.

**Table 2:** Classification Performance on (**A**) Multi-class Cancers and (**B**) Gastro-intestinal Tract Cancers.

|  |  | Predicted Class |  |  |  |  |  |  |
| --- | --- | --- | --- | --- | --- | --- | --- | --- |
|  |  | Healthy | BrCa | CRC | GBM | HBC | NSCLC | PAAD |
|  |  | Genes: 596 |  |  |  |  |  |  |
| Actual Class | Healthy | 95% | 2% |  | 2% |  | 1% |  |
|  | BrCa | 1% | 73% | 8% |  |  | 12% | 6% |
|  | CRC |  | 8% | 65% |  |  | 16% | 11% |
|  | GBM | 19% |  | 1% | 73% |  | 5% | 2% |
|  | HBC | 5% |  | 14% | 3% | 55% | 15% | 8% |
|  | NSCLC | 5% | 13% | 15% | 2% |  | 61% | 4% |
|  | PAAD |  | 6% | 13% |  | 1% | 8% | 72% |

**B**
|  |  | Predicted Class |  |  |
| --- | --- | --- | --- | --- |
|  |  | CRC | HBC | PAAD |
|  |  | Genes: 144 |  |  |
| Actual Class | CRC | 87% | 1% | 12% |
|  | HBC | 10% | 78% | 12% |
|  | PAAD | 25% | 3% | 72% |

Accuracy: 80%, Recall: 79%

**Table 3:**
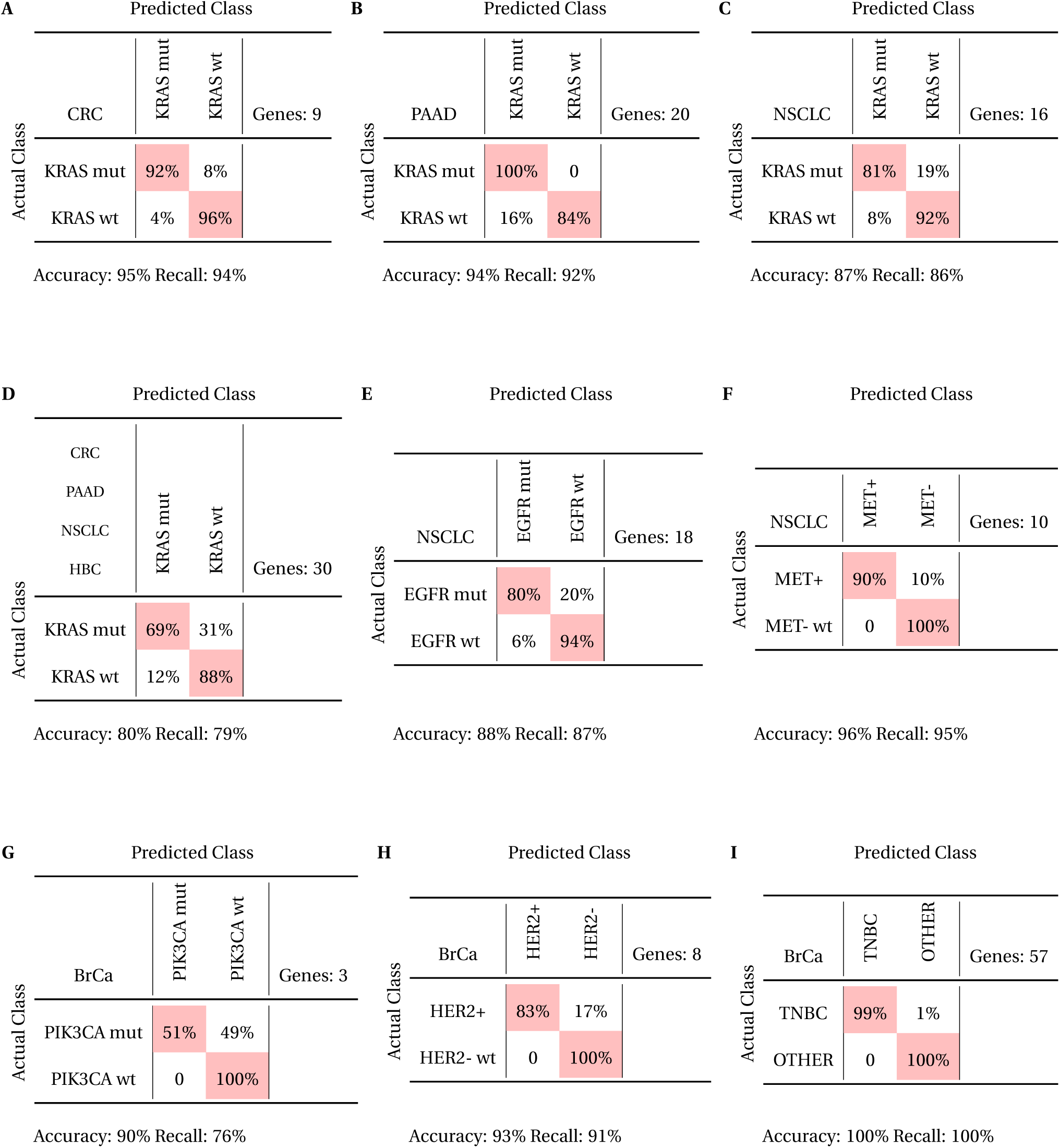
Classification Performance for Molecular Pathway Diagnostics

### Identification and analysis of biomarkers for subgrouping medulloblastoma microarray data

Medulloblastoma (MB) is the most common malignant brain tumor in children and represents approximately 20% of childhood brain tumors ^41^. Transcriptional profiling of MB identified four distinct molecular subgroups: WNT (Wnt signaling pathway), SHH (sonic hedgehog signaling pathway), Group 3 (G3) and Group 4 (G4) ^42^. Northcott *et al.* ^43^ subgrouped medulloblastoma samples by measuring the expression level of 22 subgroup-specific genes (CodeSet). Here, the DDR method was applied for identifying signature genes for each MB subgroup using microar-ray dataset (Accession: GSE37418 ^44^), and these genes were used to subgroup medulloblastoma samples from different microarray platforms. Four reference genes (*C1orf127, ZNF347, WDR70* and *HNRNPK*) (Fig. S7) were selected and DEGs (adjusted *p*-value < 1×10^−5^) for each subgroup were identified by performing DDR for each subgroup against the other subgroups (Supplementary Excel VIII). Then, top 5 genes (non-coding RNAs were not included) for each subgroup were selected based on ED values. A total of 20 genes included: WNT (*WIF1, GAD1, DKK2, TRDV3, SHOX2*), SHH (*PDLIM3, EYA1, HHIP, CRB1, SFRP1*), G3 (*TRIM58, GABRA5, PALMD, NPR3, HLX*), G4 (*EOMES, NWD2, PTPN5, RBM24, UNC5D*), and their expression heatmap is presented in Fig. 3A). Among these 20 genes, 12 overlap with medulloblastoma subgroup-specific signature genes (CodeSet) from NanoString Technologies, Inc. ^43^. The tiered catego-rizations of 20 signature genes (Supplementary Excel IX) were used as input to OneVsRestClassifier from *Scikit-learn* ^35^ using SVM over 1,000 Monte Carlo cross-validation (MCCV) iterations to yield overall accuracy of 99% and recall of 99% (Table 4A).

**Figure 3:**
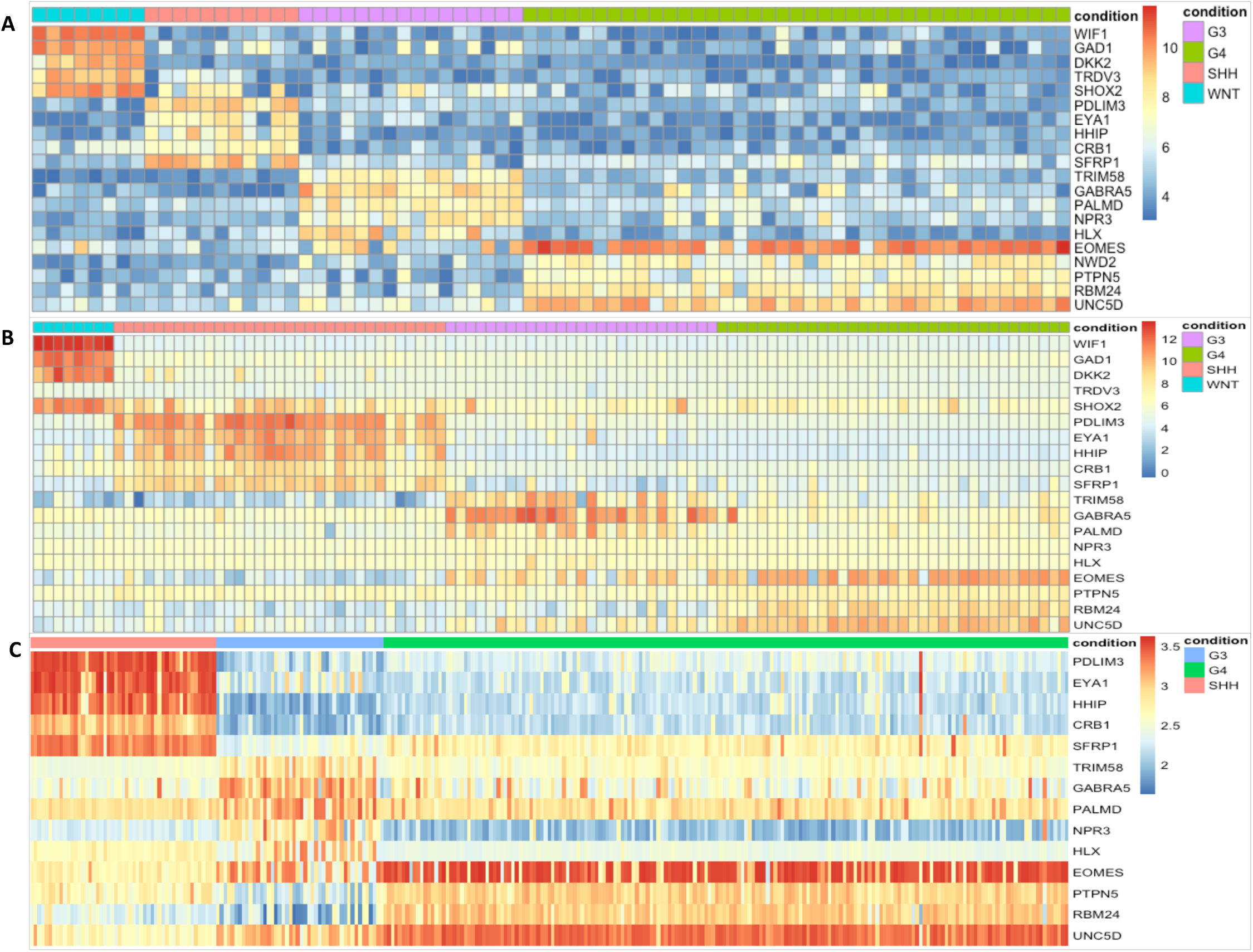
The expression heatmaps for signature genes in GSE37418^44^ (**A**), GSE21140^42^ (**B**), and GSE37382^45^ (**C**).

**Table 4:**
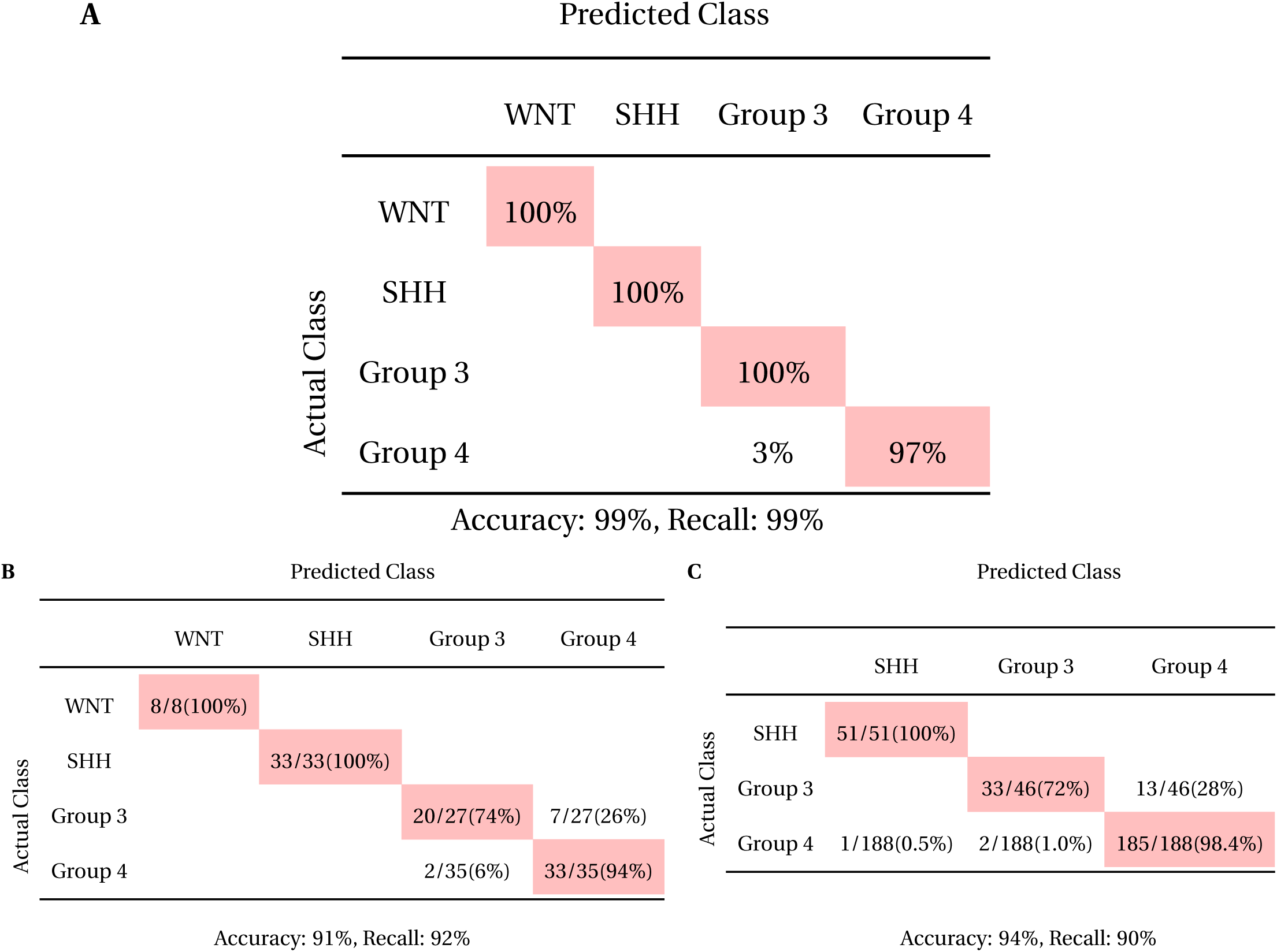
Classification Performances on medulloblastoma samples (**A**) cross-validation analysis for GSE37418 dataset, (**B**) GSE21140 dataset, and (**C**) GSE37382

The classification capacity of identified signature genes and OneVsRest classifier was evaluated on two independent datasets ((Accession: GSE21140 ^42^ and GSE37382 ^45^) from different microarray platforms. Since signature gene *NWD2* from training dataset (GSE37418) was not available in GSE21140 datasets, the other 19 signature genes (Supplementary Excel IX) were used for classification (Fig. 3B). Using OneVsRest SVM classifier trained on GSE37418 dataset, 94/103 (∼ 91%) samples from GSE21140 were assigned to appropriate subgroups (Table 4B). In GSE21140 dataset, all WNT and SHH samples were correctly classified, seven G3 samples were misclassified to G4 subgroup (7/27), and two G4 cases were misclassified to G3 group (Table 3B). There were only three subgroups of samples (SHH, G3 and G4) in GSE37382 dataset and 14 signature genes (Supplementary Excel IX and *NWD2* was not included) were used for subgrouping. Using OneVsRest SVM classifier trained on GSE37418 dataset, 51/51 (100%) SHH samples, 33/46 (72%) G3 samples and 185/188 (98%) G4 samples were correctly classified to appropriate subgroups, respectively, which resulted in accurate classification of 94% in GSE37382 dataset (Table 4C). To better characterize non-SHH/non-WNT (G3 and G4) MB, we applied DDR using the same reference genes as above to identify DEGs between G3 (adjusted *p* value < 1 × 10^−4^) and G4 (adjusted p value < 1 × 10^−5^) (Supplementary Excel VIII), then three up-regulated genes (non-coding RNAs were not included) with maximum ED values from DEGs were selected for G3 and G4, respectively (Fig. S8A). Using SVM classifier, we correctly classified G3 and G4 sub-groups with average 96% accuracy using MCCV (Table S5A). Subsequent validation using six signature genes and SVM classifier trained on GSE37418 dataset, yielded accuracies of 90% and 94% in GSE21140 and GSE37382 datasets, respectively, when subgrouping G3 and G4 (Fig. S8B and S8C). More G3 cases were correctly assigned in both validation datasets compared with predictions above (Table S5B and S5C). It is worthwhile to note that the expression level of *EN2*, which improved G3/G4 classification performance, has been reported to alter glioma cell morphology ^46^.

## Discussion

In the past decades, a wide variety of methods have been developed to identify biomarkers (feature selections) for classification of diseases using gene expression profiling. However, these approaches posed serious reproducibility challenge when classifying cross-platform samples individually due to technological and platform biases. To overcome this limitation, we present a simple, yet powerful data-driven method that does not require distribution-based modeling for gene expression analysis and it identifies potential biomarker genes with “built-in” features (categorized tiers based on reference genes) for the classification of single-sample from distinct platforms.

The huge amount of gene expression profiling data has been accumulated over the past decades and deposited in public databases such as GEO ^47^ and TCGA ^48^. These data can be great resources to detect significantly differentially expressed genes (DEGs). These DEGs may be considered as potential biomarkers for disease classification or therapeutic targets ^20^. In this study, we evaluated the ability of DDR to identify DEGs in multiple platforms through comparing with several well-established methods, *edgeR* ^32^, *DESeq* ^30^ and *DESeq2* ^31^ on two RNA-Seq datasets: TCGA-BRCA and TCGA-LUAD, and *limma* ^33^ on a microarray prostate dataset. Although not a best performer, DDR still had relatively high precision and recall values for detecting differentially expressed genes across gene expression profiling platforms. More importantly, DDR controlled the number of false positives better, which guaranteed identifying reliable biomarkers. Unlike *edgeR, DESeq, DESeq2* and *limma*, DDR is a data-driven non-parametric method which requires fewer assumption about data and is robust to outliers. As a result, it deals with cross-platform profiling gene expression and technical bias well. The utilization of reference genes at different expression levels effectively combines *p*-value and fold-change to identify reliable DEGs (biomarkers). In addition, employing logarithmic expression levels when selecting reference genes provides wiggle room to deal with overdispersion in RNA-Seq data. Our method provides a better methodological advantage to identify reliable and reproducible potential biomarkers from various expression profiling platforms. DDR can also be employed to detect DEGs among multiple conditions by designing appropriate contingency tables.

TCGA is a comprehensive molecular profiling project that compiles clinical and genomic data from samples of different human tumor types ^48^, and provide a great opportunity to identify genome-wide biomarkers for the classification of cancer conditions. In this study, we employed TCGA-BRCA datasets as training datasets and used DDR to identify small subsets of biomarkers for discriminating TNBC from the other subtypes of BRCA. The “built-in” features used as input to the classifiers effectively eliminated platform-based biases and avoided perpetuating biases from one sample into another. DDR’s ability to identify cross-platform features and classify single sample can leverage information from gene expression data that have been accumulated over the past decade and integrate them with samples now being profiled with next generation technologies. Additionally, the simplicity of these features makes them robust across various classifiers in spite of our using of SVM classifier for the classification in this study. The ability of DDR in classifying individuals into appropriate disease groups makes it an ideal choice in personalizing tool and a therapeutic strategy based on specific subgroups of cancer.

Tumor-educated platelets (TEP) is able to serve as potential noninvasive source of tumor-related RNA biomarkers ^39,40,49^. Best *et al.* showed RNA-Seq profiling of TEP could pinpoint the location of primary tumor with 71% accuracy across six types of tumors and distinguish cancer patients with different molecular subtypes as well. In this study, we applied the DDR method to identify potential biomarkers using RNA-Seq data of TEP and identify multiclass cancer and molecular subclass. We were successful in achieving comparable classification performance with fewer biomarkers compared with results reported by Best *et al.* ^39^. Much smaller sets of biomarkers associated with cancer molecular subtypes (e.g. *MET* and *HER2*-positive, or *EGFR, PIK3CA* and *KRAS* mutations) were identified by DDR as compared to biomarker sets from Best *et al.*, which makes it more practical in blood-based cancer diagnostics and therapeutic target identification. Selection of smaller subset of biomarkers can reduce over-fitting and computational complexity by removing redundant features.

DDR employs Fisher’s exact test, a non-parametric method, to analyze gene expression profiling, so it could be equally applied in the analysis of microarray data. To evaluate the performance of DDR in identification of signature genes and classification of microarray data, we applied DDR to a published microarray dataset of medulloblastoma (GSE37418) and derived 20 signature genes for medulloblastoma subgroups. Among 20 signature genes, 12 genes overlay NanoString codeset ^43^ from a commercial instrument system, which suggests our method is reliable in discovering biomarkers. The application of the classifier trained on GSE37418 dataset yielded high accuracy rate for subgrouping WNT, SHH and G4 in two independently validated dataset, confirming classification reproducibility of biomarkers identified by DDR. G3 and G4 subgroups display more similarity to each other in transcriptional profiling compared with WNT and SHH subgroups ^42^, so it is a challenge to discriminate G3 from G4. In this study, we applied DDR to identify signature genes which were differently expressed between G3 and G4, and achieved improvement in G3 assignment. A recent study suggests that non-SHH/non-WNT medulloblastoma may comprise of three subgroups rather than just G3 and G4 ^50^, which may explain the observed low accuracy for G3 and G4 subtyping.

## Conclusion

The main technical novelty of this work is the combination of data-driven reference genes with non-parametric Fisher’s exact test for discovering potential biomarkers. This not only allowed us to identify differentially expressed genes but also help to extract corresponding “built-in” features based on reference genes. One of the exciting outcomes of these “built-in” features is their reliability and reproducibility in classifying disease samples involved with technical bias and cross-platform, which allow us to analyze single sample of disease. This study has shown that DDR can be a promising tool for the identification of biomarkers for precision medicine. And some expression assay (*e.g.* quantitative PCR) based on these biomarkers and reference genes can be easily developed for diagnosis, prognosis and developing individualized treatment in the future. Finally, although this study has focused on cancer classification, it could be equally useful in classification of other diseases such as Parkinson or Alzheimer’s. In conclusion, we have developed a novel, reliable and reproducible data-driven method for identification of potential biomarkers for single-sample classification.

## Methods

### Data

#### RNA-Seq from The Cancer Genome Atlas (TCGA)

All RNA-Seq read count data from TCGA lung adenocarcinoma (LUAD) project (n=594) were retrieved using the GDC Data Transfer Tool ^51^. Both LUAD and normal data were collected, resulting in 535 cancerous and 59 normal tissue samples. All RNA-Seq read count data for breast invasive carcinoma (BRCA) project (n=1222) were downloaded from TCGA. Both cancer and normal samples were collected, resulting in 1109 BRCA and 113 normal samples. Clinical files were downloaded from TCGA data portal for all BRCA samples using GDC client tool. We identified 116 triple-negative breast cancer (TNBC) samples and 812 samples of the other subtypes based on the annotation provided in the clinical files.

#### RNA-Seq from Gene Expression Omnibus (GEO)

RPKM normalized RNA-Seq data of LUAD were downloaded from the Gene Expression Omnibus (GEO, accession: GSE40419) ^52^. The dataset contains 87 lung adenocarcinomas samples and 77 corresponding normal samples.

#### Microarray data from GEO

We retrieved mRNA expression microarray data set from GEO under the accession number GSE62872 ^53^ which was generated using platform GPL19370. These 424 samples consisting of 264 samples of prostate tumor and 160 samples of normal tissue were used. Additionally, the microarray data of 5 TNBC samples and 14 non-TNBC samples were downloaded from GEO profile data of GSE27447 ^36^.

#### RNA-Seq of Tumor-Educated Platelets

The gene expression profiles of 285 blood platelet samples were downloaded from GEO under the accession number GSE68086 ^39^. The samples consisted of breast cancer (BRCA), colorectal cancer (CRC), glioblastoma (GBM), hepatobiliary cancer (HBC), non-small cell lung cancer (NSCLC), pancreatic cancer (PAAD) and healthy donors (HD) (Table S1).

#### Microarray data of Medulloblastoma

All microarray data are downloaded from the GEO database under assession number GSE37418 ^44^, GSE21140 ^42^, and GSE37382 ^45^, respectively. The detailed number of samples for each subtype of medulloblastom from three datasets is listed in Table S2.

## Methods

### Selection of Reference Genes

Reference genes from RNA-Seq data were identified using a data-driven approach similar to that developed by Hoang et al ^54^. The normalization of RNA-Seq read counts was performed using Trimmed Mean of M-values (TMM) in edgeR ^32,55^, and then the normalized values were transformed into Counts Per Million (CPM). The CPM values for all genes in RNA-Seq datasets were used to generate two metrics across the samples, namely, the coefficient of variation (COV) and the maximum fold change (MFC). COV was calculated for each gene *i* by dividing the standard deviation (*σ_i_*) of its CPM values by the mean 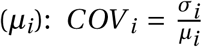. MFC is the ratio of the maximum value over the minimum CPM expression value, also for each gene. The product score (PS), our final metric for each gene, was calculated by multiplying the COV by the MFC:

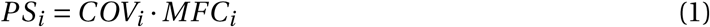

The genes with the lowest product scores and those that were also included in the list of human housekeeping genes ^56^ were selected as candidate reference genes. The list of human house-keeping genes was obtained by analyzing data from the Human BodyMap 2.0 project across 16 human tissue types ^56^. The top gene from candidate reference genes at a given range of expression was selected as the final reference gene for corresponding expression level. Let 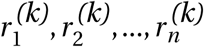 denote the expression levels of reference genes selected above in sample *k*, so that 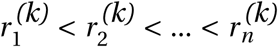. The gene expression level between 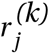 and 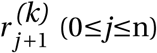 was assigned to Tier *j* in sample *k*, here 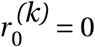 and 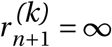 (Fig. 1).

### Identification of Discriminant Genes

A *C* ×*T* contingency table was created for each gene, here *C* is the number of conditions and *T* is the number of tiers, to display the numbers of samples from a particular condition in which that gene was assigned to a particular tier (Fig. 1). Then, Fisher’s exact tests (FETs) were performed for the contingency tables to assess whether the expression level of the gene is independent or correlated with conditions or phenotypes. The resulting *p*-values were adjusted to account for multiple tests using the p.adjust function in R (method = ‘fdr’). In addition to adjusted *p*-values, we defined expression distance (ED) for each gene to describe a quantity change of the gene expression between conditions. The ED can be used to select up- and down-regulated genes.

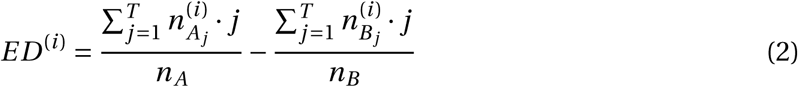

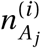 = the number of samples from group *A* with gene *i* in tier *j*

*n*_*A*_ = the number of samples from group *A*

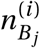 = the number of samples from group *B* with gene *i* in tier *j*

*n*_*B*_ = the number of samples from group *B*

### Benchmarks Comparison in Identification of Discriminant Genes

To assess how well DDR performs for identification of DEGs in comparison to the current methods (*DESeq* ^30^, *DESeq2* ^31^ and *EdgeR* ^32^ for RNA-Seq data and *limma* ^33^ for microarray data), we used some real and simulated data. To compute the true positives, we randomly selected the subsets with different sizes (equal number for each condition) from full datasets, and the random selection was repeated 10 times to avoid sample selection bias. Larger sample size generally leads to increased precision, so the overlapped DEGs identified from full dataset and subset approximate the set of true positives. The tested methods were compared under precision and recall ^57^ defined as

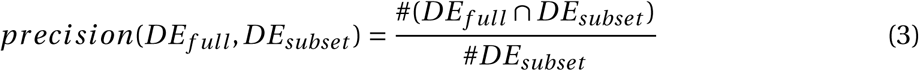

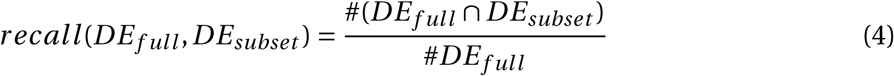

*DE*_*full*_ = the set of identified DEGs in the full data set

*DE*_*subset*_ = the set of identified DEGs in a subset of the data

To evaluate the false discovery rate of different methods, we randomly assigned the equal sizes of samples from the same condition without replacement into two groups and the procedure was repeated 10 times. All samples were from the same condition, which means that there should not be any real DEGs, so DEGs identified from simulated datasets arise by chance alone. The false discovery rate was defined as ratio of the number of identified DEGs from simulated dataset to the number of identified DEGs by comparing two conditions from complete dataset. We compared the overlaps of the identified DEGs between the methods by Szymkiewicz-Simpson overlap coefficient.

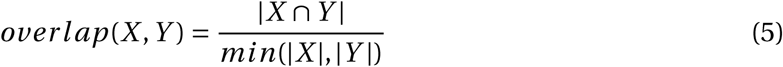

Here, *X* and *Y* are the lists of DEGs identified by two methods, respectively.

### Feature Selection and Classification

Small subset of biomarkers for classification was obtained from DEGs by filtering on the basis of low adjusted *p*-values (FDRs) and high EDs. The feature tables consisting of tiers to which selected biomarkers belong for samples served as input to the classifiers. An example of feature table for classifying TNBC and non-TNBC is shown in Table S3.

Several machine learning classifiers from *Scikit-learn* ^35^ Python library were applied to classification. We selected Support Vector Machine (SVM) using RBF kernel with C=1 as final classification model for binary classification and OneVsOneClassifier (or OnevsRestclassifier) using SVM to deal with multi-class classification problems. The samples were randomly separated into training/testing sets with 90% of samples as training and 10% as testing. And then, we followed 1,000 iterations in stratified cross-validation analysis which deals with imbalanced classes and used accuracy, precision, recall and F1 scores from *Scikit-learn* to assess the performance of our classification model.

## Supporting information

Supplementary Excel III

Supplementary Excel VII

Supplementary Excel VIII

