## Supplemental Tables and Figures for "Multiplatform Biomarker Identification using a Data-driven Approach Enables Single-sample Classification"

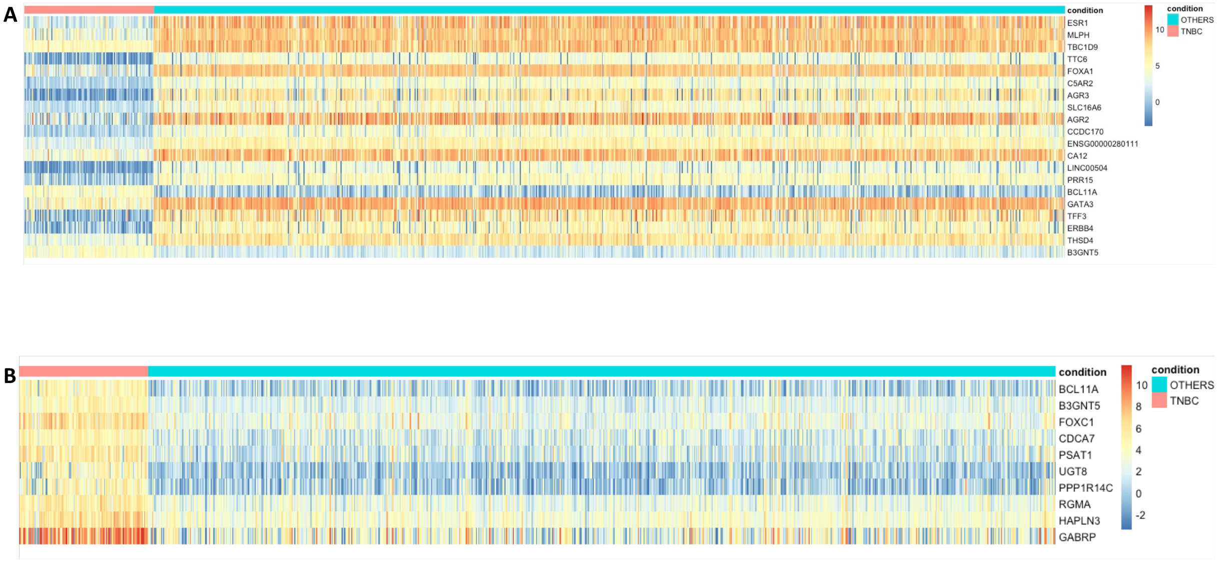


Figure S1: Expression heatmap of top 20 differentially expressed genes between triple-negative breast caner samples (red) and other types of breast cancer samples (turquoise) (**A**) and expression heatmap of top 10 up-regulated genes in triple-negative breast caner samples (red) compared with other types of breast cancer samples (turquoise) (**B**) in TCGA-BRCA RNA-Seq dataset. Expression level of gene is represented as log2(counts+1).

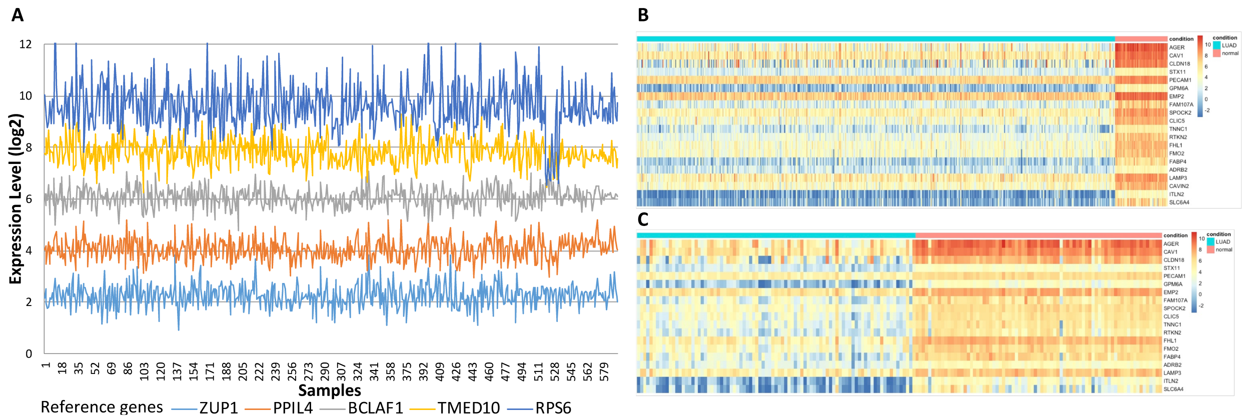


Figure S2: Expression levels of five data-driven reference genes (**A**) and expression heatmap showing top 20 differentially expressed genes between LUAD samples (turquoise) and normal samples (red) (**B**) in TCGA-LUAD RNA-Seq dataset . Expression heatmap of 19 top genes identified from TCGA-LUAD dataset between LUAD samples (turquoise) and normal samples (red) in independent validated dataset (Accession: GSE40419)(**C**).

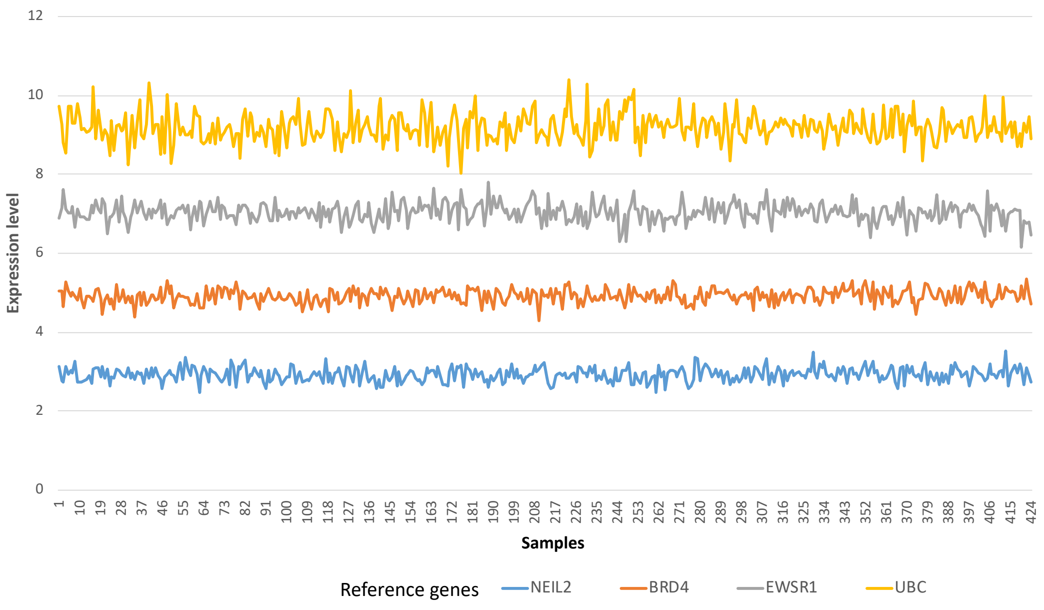


Figure S3: Expression levels of four data-driven reference genes from prostate tumor microarray samples downloaded from GEO (Accession: GSE62872).

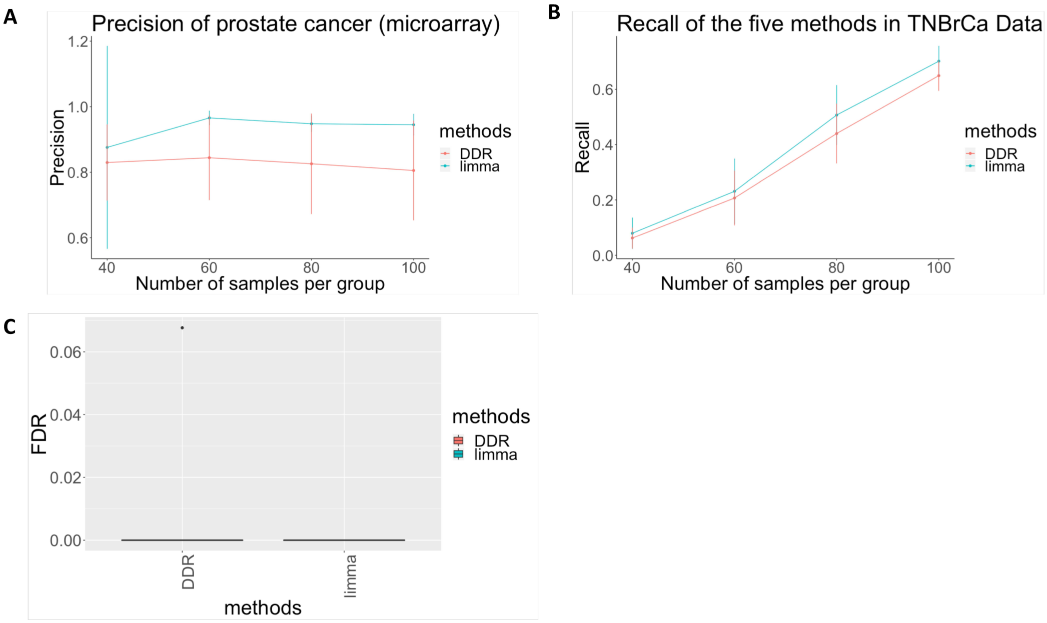


Figure S4: Precision (**A**), recall (**B**) and FPR (**C**) of methods in prostate tumor microarray dataset from GEO (Accession: GSE62872).

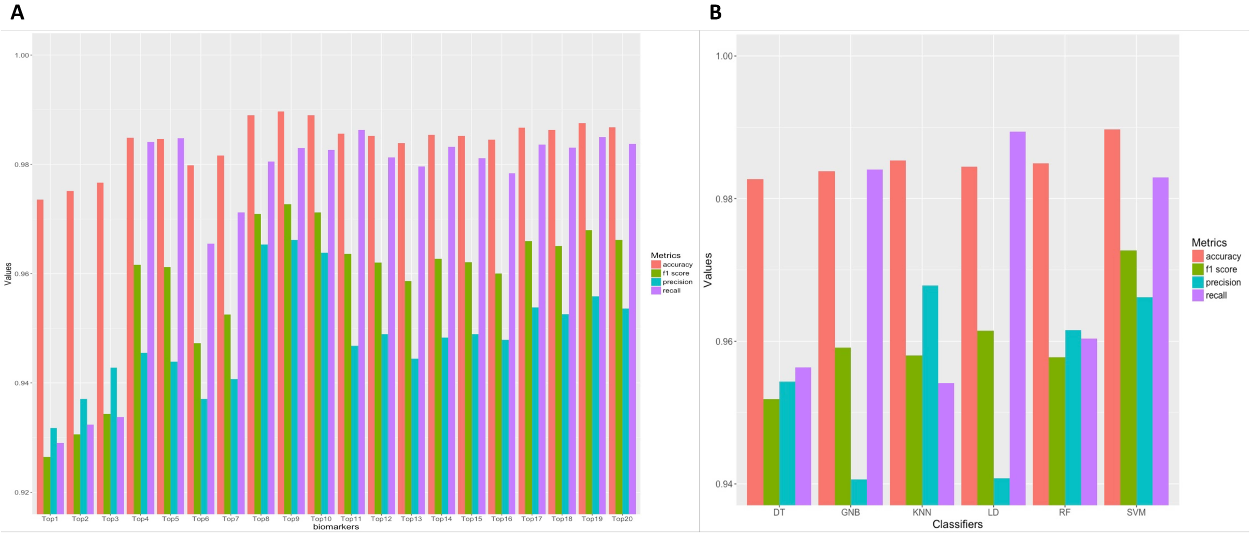


Figure S5: Classification performance of different numbers of top genes using SVM classifier (**A**) and different machine learning classifiers using classified tiers of top 9 genes (**B**) in TCGA LUAD RNA-Seq dataset.

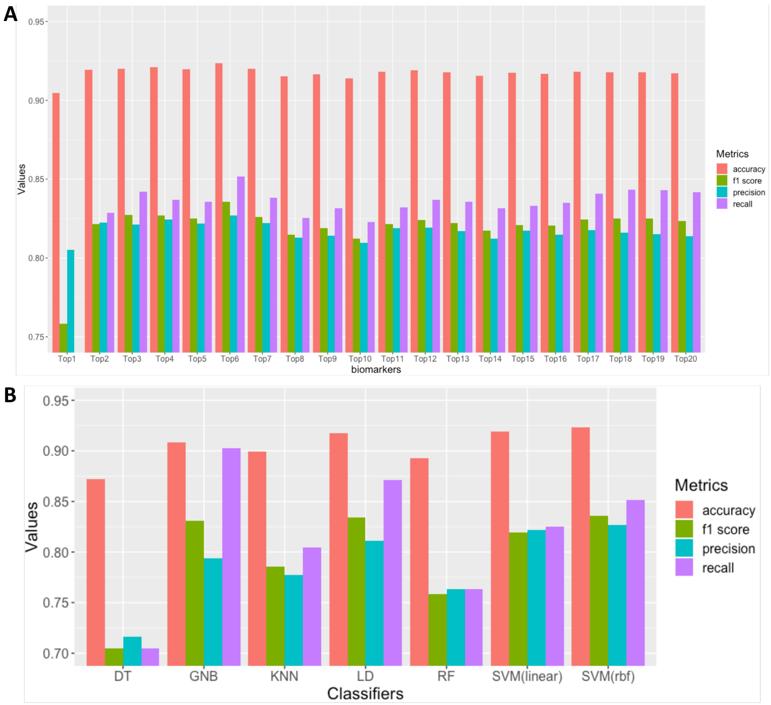


Figure S6: Classification performance of different numbers of top genes using SVM classifier (**A**) and different machine learning classifiers using classified tiers of top 6 genes (**B**) for classifying TNBC and non-TNBC in TCGA-BRCA RNA-Seq dataset.

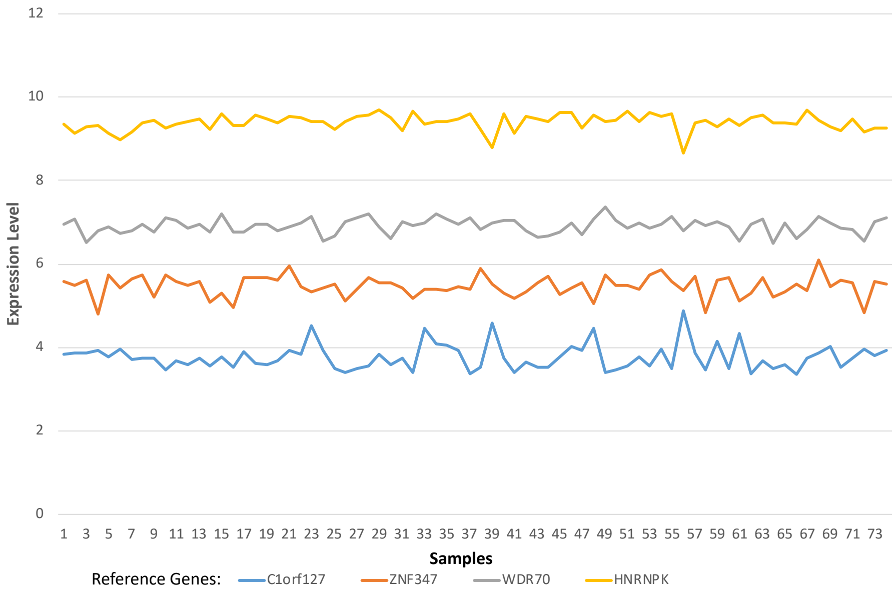


Figure S7: Expression levels of four data-driven reference genes from medulloblastoma microarray dataset from GEO (Accession: GSE37418).

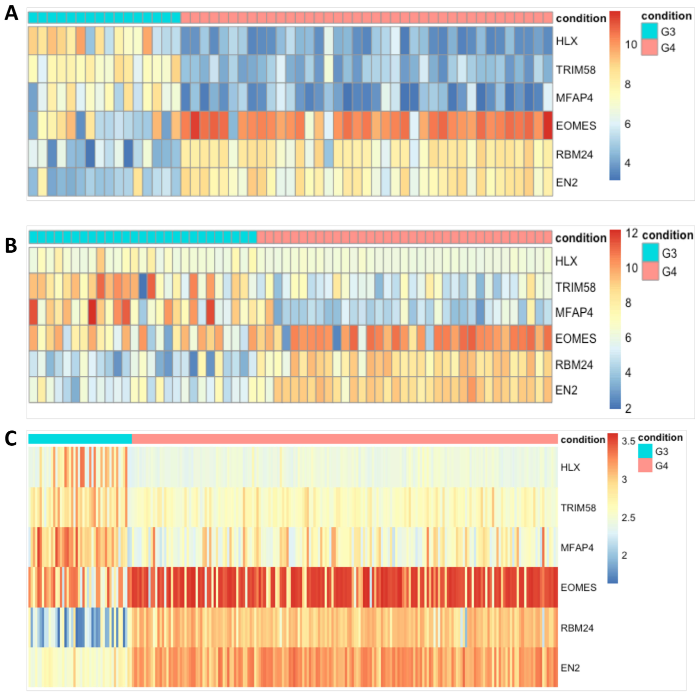


Figure S8: The expression heatmaps for signature genes between G3 (turquoise) and G4 (red) in GSE37418(**A**), GSE21140(**B**), and GSE37382(**C**).

Table S1: RNA-seq samples from Tumor-Educated Platelets

| Cancer subtype | Number of samples |
|---|---|
| Breast cancer | 39 |
| Colorectal cancer | 42 |
| Glioblastoma | 40 |
| Hepatobiliary cancer | 14 |
| Lung cancer | 60 |
| Pancreatic cancer | 35 |
| Healthy contro | 55 |

Table S2: Microarray samples for Medulloblastoma

| Subtype | GSE37418 | GSE21140 | GSE37382 |
|---|---|---|---|
| WNT | 8 | 8 | 0 |
| SHH | 11 | 33 | 51 |
| G3 | 16 | 27 | 46 |
| G4 | 39 | 35 | 188 |

Table S3: Feature table for classifying TNBC and non-TNBC

| BCL11A | B3GNT5 | Conditions |
|---|---|---|
| 2 | 2 | TNBC |
| 1 | 2 | TNBC |
| 2 | 2 | TNBC |
| 2 | 2 | TNBC |
| 0 | 0 | non-TNBC |
| 0 | 0 | non-TNBC |
| 0 | 1 | non-TNBC |
| 1 | 1 | non-TNBC |

Table S4: Overlaps of DEGs identified between the different methods in TCGA-BRCA dataset and TCGA-LUAD dataset

| **Method** | DDR | EdgeR_GLM | EdgeR_EXACT | DESeq | DESeq2 |
|---|---|---|---|---|---|
| *TCGA-BRCA dataset* | | | | | |
| DDR | 100 | 87 | 88 | 48 | 87 |
| EdgeR_GLM | | 100 | 99 | 85 | 85 |
| EdgeR_EXACT | | | 100 | 85 | 85 |
| DESeq | | | | 100 | 98 |
| DESeq2 | | | | | 100 |
| *TCGA-LUAD dataset* | | | | | |
| DDR | 100 | 80 | 80 | 52 | 80 |
| EdgeR_GLM | | 100 | 99 | 81 | 92 |
| EdgeR_EXACT | | | 100 | 81 | 92 |
| DESeq | | | | 100 | 98 |
| DESeq2 | | | | | 100 |

Table S5: Classification Performances between Group 3 and Group 4 on (A) cross-validation analysis for GSE37418 dataset, (B) GSE21140 dataset, and (C) GSE37382

**A**

| Actual Class | Predicted Class: Group 3 | Predicted Class: Group 4 |
|---|---|---|
| Group 3 | 94% | 6% |
| Group 4 | 2% | 98% |

Accuracy: 96%, Recall: 96%

**B**

| Actual Class | Predicted Class: Group 3 | Predicted Class: Group 4 |
|---|---|---|
| Group 3 | 23/27(85%) | 4/27(15%) |
| Group 4 | 2/35(6%) | 33/35(94%) |

Accuracy: 90%, Recall: 90%

**C**

| Actual Class | Predicted Class: Group 3 | Predicted Class: Group 4 |
|---|---|---|
| Group 3 | 36/46(78%) | 10/46(22%) |
| Group 4 | 5/188(3%) | 183/188(97%) |

Accuracy: 94%, Recall: 88%
